# Defocus and magnification dependent variation of TEM image astigmatism

**DOI:** 10.1101/138255

**Authors:** Rui Yan, Kunpeng Li, Wen Jiang

**Affiliations:** Markey Center for Structural Biology, Department of Biological Sciences, Purdue University, West Lafayette, IN 47907, USA; Department of Chemistry, Purdue University, West Lafayette, IN 47907, USA; Purdue Institute of Inflammation, Immunology and Infectious Disease, Purdue University, West Lafayette, IN 47907, USA

**Keywords:** defocus-dependent astigmatism, magnification-dependent astigmatism, objective lens current, octupole stigmators, vector summation, cryo-EM

## Abstract

Daily alignment of the microscope is a prerequisite to reaching optimal illumination lens and imaging lens conditions for high resolution imaging in cryo-EM. In contrast to the dramatic progress in automated image acquisition and post-image processing techniques, less attention has been paid to the improvement of microscope alignment before data collection. In this study, we have employed our recently published tool, *s^2^stigmator*, to study how image astigmatism varies with the imaging conditions (e.g. defocus, magnification). We have found that the large change of defocus/magnification between visual correction of astigmatism and subsequent data collection tasks, or even during data collection will inevitably result in undesirable residual astigmatism in the final images. Furthermore, the dependence of astigmatism on the imaging conditions varies significantly from time to time, so that it cannot be reliably compensated by pre-calibration of the microscope. These findings have essentially invalidated a basic assumption of current cryo-EM imaging strategies that assumes invariant astigmatism for different defocuses/magnifications used in the microscope alignment stage and the final data acquisition stage. Based on these findings, we recommend the same magnification and the median defocus of the intended defocus range for final data collection are used in the objective lens astigmatism correction task during microscope alignment and in the focus mode of the iterative low-dose imaging. It is also desirable to develop a fast, accurate method that can perform dynamic correction of the astigmatism for different intended defocuses during automated imaging.

## 1. Introduction

Cryo-electron microscopy (cryo-EM) has become a powerful technique for structural studies of macromolecular complexes and assemblies at near-atomic resolutions. Good alignment of the microscope, such as the gun, condensers, apertures, beam tilt, coma (Ishizuka, 1994; Zemlin et al., 1978), astigmatism, *etc.*, is a prerequisite to reaching optimal illumination lens and imaging lens conditions for high resolution TEM imaging. Currently, microscope alignments are still performed manually and rely on visual, qualitative feedback. Moreover, manual microscope alignment requires extensive training and experience but still often suffers from suboptimal efficiency and quality. Minimizing astigmatism of the objective lens is an indispensable daily instrument alignment task essential for high resolution TEM imaging.

Astigmatism of the objective lens represents the angular dependency of defocus. 2-fold astigmatism is the major type of astigmatism relevant to cryo-EM, which results in the elliptical elongation of Thon rings (Thon, 1971) in the power spectra of TEM images. Currently, many microscopists follow a common approach (Grassucci et al., 2008; Sun and Li, 2010) which is to visually correct astigmatism at large magnifications and small defocuses, then switch to a drastically different imaging condition to collect data by intentionally varying defocus to sample the entire reciprocal space and even-out the zero-nodes of the contrast transfer function (CTF) (Cheng et al., 2015; Penczek, 2010; Zhu et al., 1997). The implicit assumption for this common strategy is that the astigmatism is invariant to the change of magnification and defocus. However, such invariance has not been quantitatively validated, and on the contrary, we have now shown in this study that the assumption is incorrect.

Due to the poor sensitivity of human eyes, microscopists have to rely on large magnifications and small defocuses when they visually examine the roundness of Thon rings in the 2D power spectra displayed on a computer screen and iteratively adjust the two objective lens stigmators to make the Thon rings as circular as possible. This tedious and subjective method is not only inaccurate and potentially biased by the astigmatism of human eyes, but also hampers the systematic quantification of the astigmatism variations for different magnifications/defocuses. To overcome this challenge, it is desirable to take advantage of a proper approach which is able to sensitively measure and accurately correct astigmatism at any imaging condition.

We have recently published a method, *s^2^stigmator* (Yan et al., 2017), with a single-pass tuning strategy, that allows rapid and sensitive detection of astigmatism using TEM live images and can reliably and efficiently guide the user to manually adjust the two stigmators to correct astigmatism. This exciting method opens up possibilities to minimize astigmatism with real-time feedback at a wide range of imaging conditions that are not available by visual examination. In this article, we present systematic and quantitative investigations of astigmatism dependence on imaging conditions by employing *s^2^stigmator* to correct astigmatism and then varying imaging conditions (defocus, magnification). Underlying physical principles are used to interpret the variability of astigmatism and its dependence on image conditions. Based on the findings of these studies, several recommendations are provided for instrument alignment and data acquisition to help maximally reduce astigmatism and improve high resolution imaging.

## 2. Method

### 2.1 Experimental cryo-EM datasets for initial test

Our study started with three datasets of experimental cryo-EM micrographs which were downloaded from EMPIAR (Iudin et al., 2016) or collected by our own group. The MAVS CARD filament dataset (EMPIAR-10014), plasmodium falciparum 80S ribosome dataset (EMPIAR-10028), and our own RNA polymerase dataset (unpublished) were acquired using a JOEL 2200FS, a FEI POLARA and a FEI Titan Krios microscope at a nominal magnification of 60,000X, 78,000X and 22,500X, respectively. For each dataset, defocus and astigmatism were estimated using *ctffind3* (Mindell and Grigorieff, 2003) in order to examine the correlation between them.

### 2.2 Data collection for the study of defocus-dependent astigmatism

Next, the defocus-dependence of astigmatism was examined using live images of carbon film obtained on our CM200 microscope at 200 kV and FEI Titan Krios at 300 kV, and recorded on a Gatan US4000 CCD and K2 Summit camera, respectively. We minimized the objective lens astigmatism at small, medium and large defocuses using our *s^2^stigmator* tool, then increased or decreased defocus from the starting defocus used for astigmatism correction. At each defocus, ten images were collected and their defocuses, astigmatisms were calculated to obtain their mean and root-mean-square deviation (RMSD) values. The same experiment was repeated on different days to explore the reproducibility of defocus-dependence of astigmatism.

### 2.3 Data collection for the study of magnification-dependent astigmatism

Finally, the magnification-dependence of astigmatism was examined using the same sample and instrument as described above. In order to emulate the change of magnification between astigmatism correction and data collection, we minimized the astigmatism at a high magnification using *s^2^stigmator*, then successively lowered the magnifications. At each magnification, twenty images were collected, and their mean and RMSD of astigmatism and defocus were computed. In addition to the magnification-dependence of astigmatism, magnification-dependence of defocus was also simultaneously examined using the same set of data. The same measurement was also repeated multiple times in order to detect the stability of the relationship between astigmatism and magnification.

The magnifications were calibrated using polycrystalline gold sample grids. The anisotropic magnification distortion (Grant and Grigorieff, 2015; Yu et al., 2016) was corrected from the live images according to the previously determined parameters before the astigmatism was calculated.

## 3. Results

### 3.1 Observations of defocus-dependent astigmatism in experimental cryo-EM data

To examine the defocus-dependence of objective lens astigmatism, we first calculated the defocus and astigmatism of three cryo-EM datasets as described in Section 2.1 and plotted the results in Fig. 1. It can be seen that there are positive correlations between astigmatism and defocus, providing evidence for defocus-dependent astigmatism in experimental cryo-EM data. It is worth pointing out that this correlation is a general phenomenon since it is observed in a wide variety of data, such as data from multiple research groups, different vendors’ instruments, a diversity of samples, and varying imaging conditions. In our own Titan Krios data (Fig. 1C), the astigmatism was calculated after correction of anisotropic magnification distortion (Grant and Grigorieff, 2015; Yu et al., 2016). The anisotropic magnification distortion in the other two EMPIAR (Iudin et al., 2016) datasets (Fig. 1A, B) was not corrected as the distortion parameters were not known. However, these two datasets were both imaged at high magnifications in which distortion is generally negligible. As illustrated in Fig. 1, the variations of astigmatism for different defocuses are significant (e.g. >100 nm) and distinct from dataset to dataset. Hence, it is desirable to comprehensively examine the dependence of astigmatism on imaging conditions (e.g. defocus, magnification) which are frequently changed in TEM alignment and during data acquisition.

**Fig. 1.**
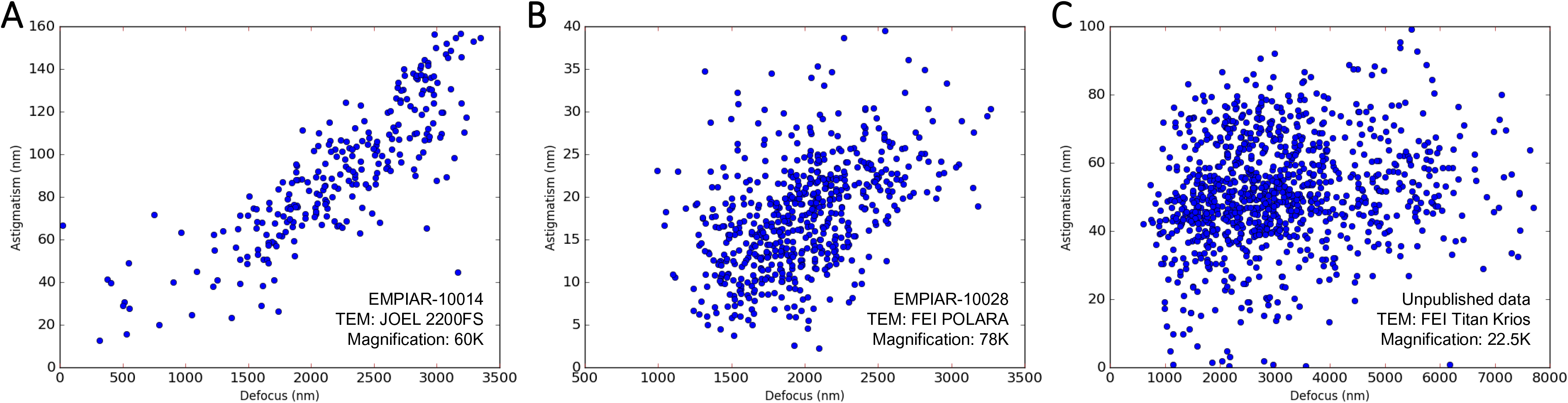
Observations of the relationship between defocus and astigmatism in experimental cryo-EM datasets. The EMPIAR ID (Iudin et al., 2016), instrument, and magnification are marked at the lower right corner of each plot.

### 3.2 Robust performance of *s^2^stigmator* and the single-pass tuning strategy at different defocuses and magnifications

For quantitative measurement of astigmatism variations, astigmatism needs to be accurately corrected at a wide range of imaging conditions, which is very challenging for the current method relying on visual examination. We have recently published a closed-form algorithm, *s^2^stigmator*, with a single-pass tuning strategy (Yan et al., 2017), that allows fast and sensitive detection of astigmatism using TEM live images and guides the users to reliably and efficiently adjust the two objective lens stigmators to correct astigmatism. Fig. 2 displays two screenshots of the entire trajectories acquired from astigmatism correction processes on Titan Krios (Fig. 2A) and CM200 microscopes (Fig. 2B), respectively. The gray level of the points is varied to make more recent ones darker in order to clearly display the sequence of the points. The user would first adjust either of the two stigmators to find the optimal value (points marked by blue arrows in Fig. 2) that gives rise to minimal astigmatism in the arc-shaped trajectory; then adjust the other stigmator to linearly move the points to the center at which the astigmatism is 0. Thus, *s^2^stigmator* was used in this study for real-time determination and minimization of astigmatism for systematic studies of objective lens astigmatism.

**Fig. 2.**
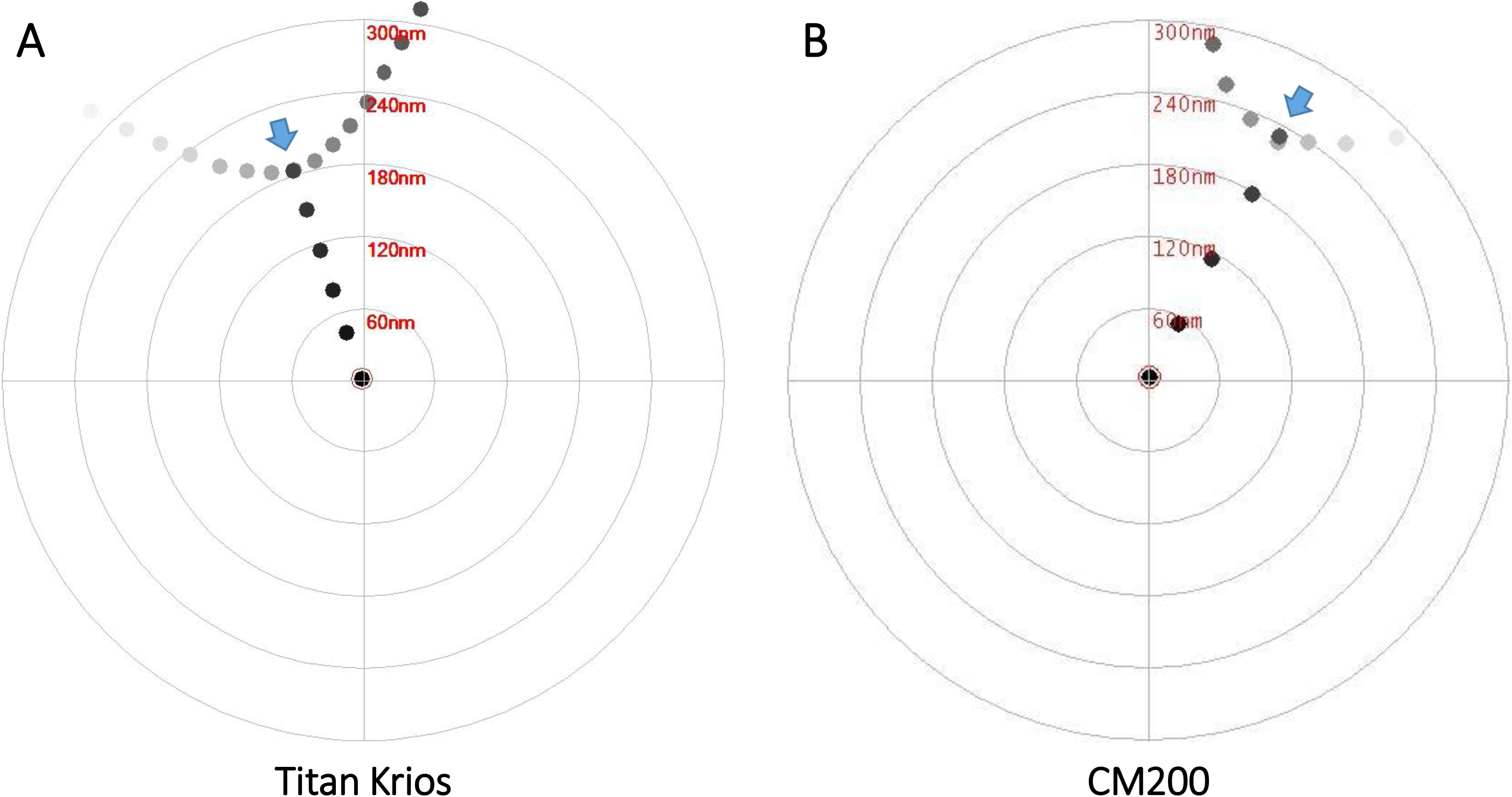
Performance of *s^2^stigmator* method and the single-pass tuning strategy. (A) A screenshot of the trajectory from Titan Krios microscope obtained at 1000 nm defocus, a nominal magnification of 22,500X and 25 e/Å^2^ dose with images recorded on a Gatan K2 Summit direct electron detector operated at counted mode using 15 e/pixel/second dose rate and 3s exposure time. Stigmator MX was adjusted first (arc-like segment) and then MY was adjusted (straight segment). (B) A screenshot of trajectory from CM200 microscope obtained at 1000 nm defocus, a nominal magnification of 66,000X and 40 e/Å^2^ dose with images recorded on a Gatan UltraScan 4k CCD with 3s exposure time. Stigmator MY was adjusted first (arc-like segment) and then MX was adjusted (straight segment). The wide blue arrow indicates the optimal point with minimum astigmatism in the arc-like segment of each trajectory.

Next, we tested the performance of our *s^2^stigmator* method by correcting astigmatism at various imaging conditions, including defocus and magnification. Fig. S1 shows the screenshots of trajectories obtained from Titan Krios when correcting astigmatism at different defocuses (Fig. S1A-C) and different magnifications (Fig. S1D-F). In these six screenshots, the trajectories are very similar and all consistently led to correction of astigmatism at a wide range of defocuses and magnifications. The angle of the straight trace segment corresponds to the 2nd stigmator used in this single-pass strategy, i.e. stigmator MX in Fig. S1. This angle is determined by the angular position of the stigmators, e.g. octupole objective lens stigmator (Hawkes, 2013; Rai-Choudhury, 1997). Fig. S2 shows the trajectories acquired at a variety of defocuses (Fig. S2A-C) and magnifications (Fig. S2D-F) on CM200. Similarly, the change of defocus on CM200 in the range of cryo-EM research does not have a significant influence on the shape of the trajectories (Fig. S2A-C). However, the switch of magnification does have an effect on the orientation of the trajectories (Fig. S2D-F). It is noted that the trajectories turn clockwise when magnification increases (Fig. S2D-F), which is consistent with the rotation of real images at the same set of magnifications (Fig. S2G-I). Moreover, the angle changes of the straight trace segments between two adjacent magnifications are 14° (65° in Fig. S2D v.s. 51° in S2E) and 26° (51° in Fig. S2E v.s. 25° in S2F), identical to the change of angles (marked by red dash lines in Fig. S2G-I) among the corresponding real images (14° between Fig. S2G and S2H, 26° between Fig. S2H and S2I). Thus, we attribute the rotation of trajectories at different magnifications to the imperfect implementation of the rotation-free imaging function on CM200. In contrast, the rotation-free imaging function on Titan Krios is excellent as shown by the absence of rotations of the trajectories in Fig. S1D-F for different magnifications.

### 3.3 Defocus-dependent astigmatism

After confirming that the astigmatism of objective lens could be accurately minimized using *s^2^stigmator*, we systematically investigated the dependence of astigmatism on defocus using live images of carbon film at room temperature. We first corrected the astigmatism at a specific defocus, then measured the astigmatism with all other instrument parameters remaining constant while only the defocus was gradually altered with a fixed step size (e.g. 100 nm). Fig. 3 shows clear correlation between defocus and astigmatism for both Titan Krios (Fig. 3A-D) and CM200 (Fig. 3E-H) microscopes at a nominal magnification of 22,500X and 115,000X, respectively. At each defocus, the point and error bar represent the mean and RMSD of astigmatism from ten images, respectively. When the astigmatism is minimized at small, medium, and large defocus (red, green and blue lines in Fig. 3A, E), astigmatism linearly increases as the defocus is continuously increased/decreased from the starting defocus used for astigmatism correction. Apparently, the slopes of the lines from CM200 (Fig. 3E) are much larger than those from Titan Krios (Fig. 3A), implying the dependence of astigmatism on defocus for CM200 is much more severe than that for Titan Krios. Furthermore, polar plots were used to show the raw data distribution of the astigmatism used for the line graphs with the same colors. Fig. 3B-D present the polar distribution of astigmatism with varying defocus when the astigmatism is corrected at small (Fig. 3B), medium (Fig. 3C), and large defocus (Fig. 3D) on Titan Krios, corresponding to the red, green, and blue line in Fig. 3A, respectively. It is evident that the astigmatism angle stably points to a certain direction as the astigmatism amplitude gradually increases due to the monotonically ascending (Fig. 3B) or descending defocus (Fig. 3D). In addition, Fig. 3C shows that the astigmatism angle changes about 90° when defocus changes bi-directionally after astigmatism minimization. The 90° angle change corresponds to the swapping of the major and minor axes of astigmatism. Similar distributions can also be seen from the raw data acquired on CM200 (Fig. 3F-H) but with larger increase of astigmatism than that on Titan Krios (Fig. 3B-D). Therefore, all the analyses above demonstrate the general existence of defocus-dependent astigmatism in the objective lens of TEM.

**Fig. 3.**
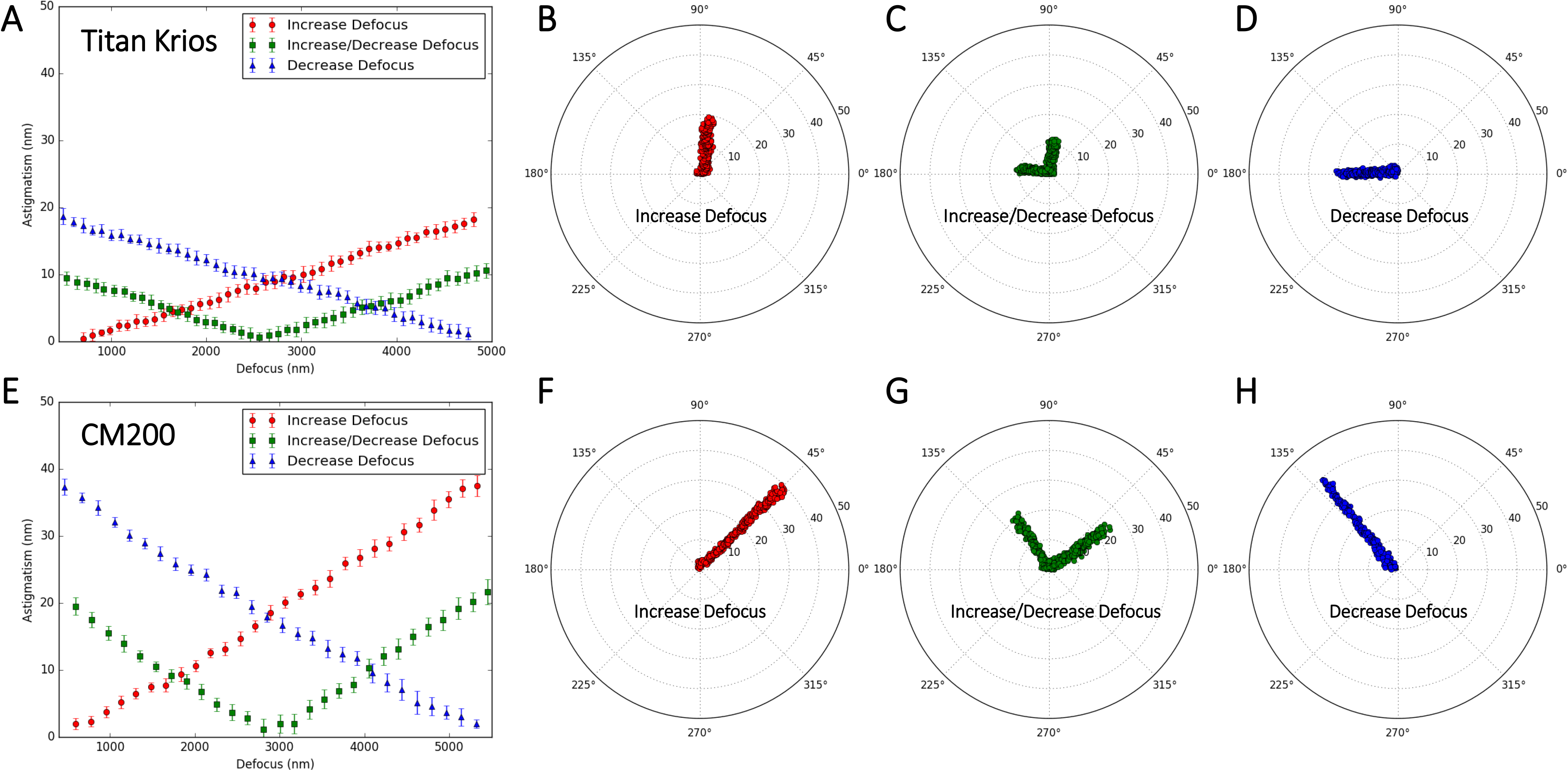
Defocus-dependent astigmatism. (A) The increment of astigmatism with the change of defocus on Titan Krios microscope when the astigmatism is corrected at small (red), medium (green) and large (blue) defocus, respectively. (B-D) The polar distribution of all data obtained from Titan Krios microscope when the astigmatism is corrected at small (B), medium (C) and large (D) defocus, corresponding to the line with the same color in (A), respectively. (E) The increment of astigmatism with the change of defocus on CM200 microscope obtained from the same experiment described in (A). The correlation between (E) and (F-H) is the same as that between (A) and (B-D). In the line plots (A, E), the point and error bar at each defocus represent the mean and RMSD of astigmatism from ten images, respectively.

For a comprehensive understanding of defocus-dependent astigmatism, we repeated our measurement on different days and compared the variation of astigmatism. As shown in Fig. 4A and E, the slopes of lines are not identical even for the measurements made using the same conditions, demonstrating the amount of dependence varies from day to day on both Titan Krios (Fig. 4A) and CM200 (Fig. 4E) microscopes. What's more, much more pronounced differences in the astigmatism angles were shown in the data collected on Titan Krios (Fig. 4B-D) among different days when the astigmatism is initially minimized at a small defocus. The differences indicate that the distribution of defocus-dependent astigmatism cannot be exactly reproduced even though the defocus is adjusted in the same way. This irreproducibility can also be observed from the data collected on CM200 (Fig. 4F-H) when the astigmatism correction is performed at a large defocus. Consequently, the comparison of the repetitive measurements confirms the variability of defocus-dependent astigmatism in the objective lens of TEM.

**Fig. 4.**
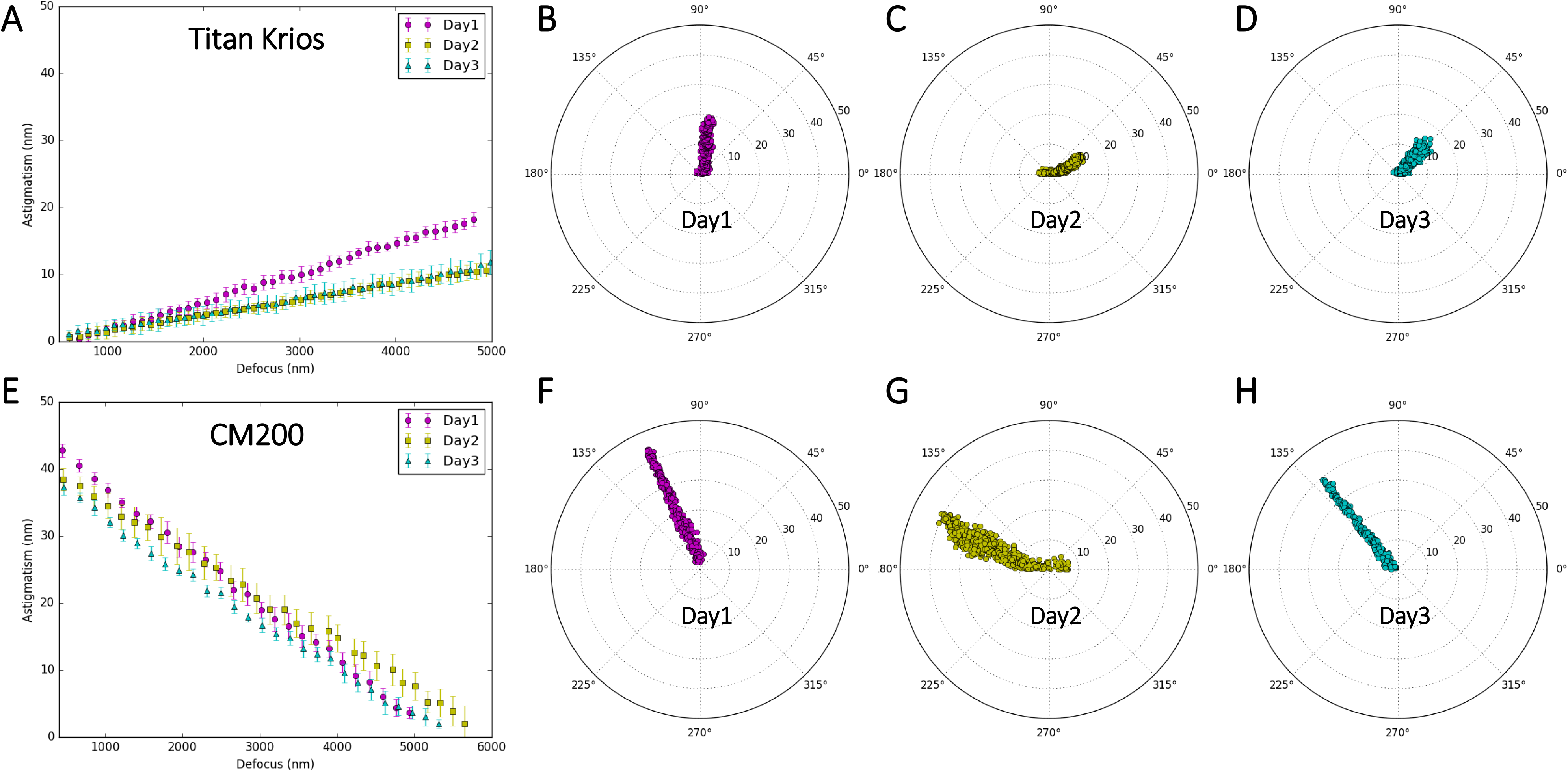
Variability of defocus-dependent astigmatism. (A) The profile of astigmatism increment as defocus increases on Titan Krios microscope when the astigmatism is minimized at small defocus on three different days. (B-D) The polar distribution of all data from the repeated experiments on Titan Krios microscope described in (A). Each polar distribution corresponds to the line with the same color as in (A). (E) The profile of astigmatism increment as defocus decreases on CM200 microscope when the astigmatism is minimized at large defocus on three different days. The correlation between (E) and (F-H) is the same as that between (A) and (B-D). In the line plots (A, E), the point and error bar at each defocus represents the mean and RMSD of astigmatism from ten images, respectively.

### 3.4 Magnification-dependent astigmatism

We also used *s^2^stigmator* to investigate the dependence of astigmatism on magnification and to test the implicit assumption of invariant astigmatism at different magnifications for the common practice of using a higher magnification for correction of astigmatism than that for data acquisition. We first corrected astigmatism at a nominal magnification of 96,000X on Titan Krios and 250,000X on CM200 microscope, respectively, then measured the astigmatism as the magnification was stepwise reduced while keeping all other instrument parameters unchanged. Fig. 5 shows the change of astigmatism/defocus with magnification for Titan Krios and the variability of this change across multiple measurements. At each magnification, we collected twenty images, plotted the distribution of astigmatism in polar coordinate (Fig. 5A-C), and then calculated their mean and RMSD of astigmatism (blue line)/defocus (red line) represented as points and error bars (may be too small to be visible) in Fig. 5D-F. When repeating the measurements on Titan Krios, the changes of defocus (red lines in Fig. 5D-F) follow a similar pattern, but the profiles of both astigmatism amplitude (blues lines in Fig. 5D-F) and astigmatism angle (Fig. 5A-C) are irreproducible. Similar results were also found for CM200 (Fig. 6) in which the defocus tends to increase with lower magnifications (red lines in Fig. 6D-F), rather than decrease as shown for Titan Krios (red lines in Fig. 5D-F). All these data demonstrate the existence of magnification-dependent astigmatism and its stochastic fluctuations for both high-end and low-end TEM.

**Fig. 5.**
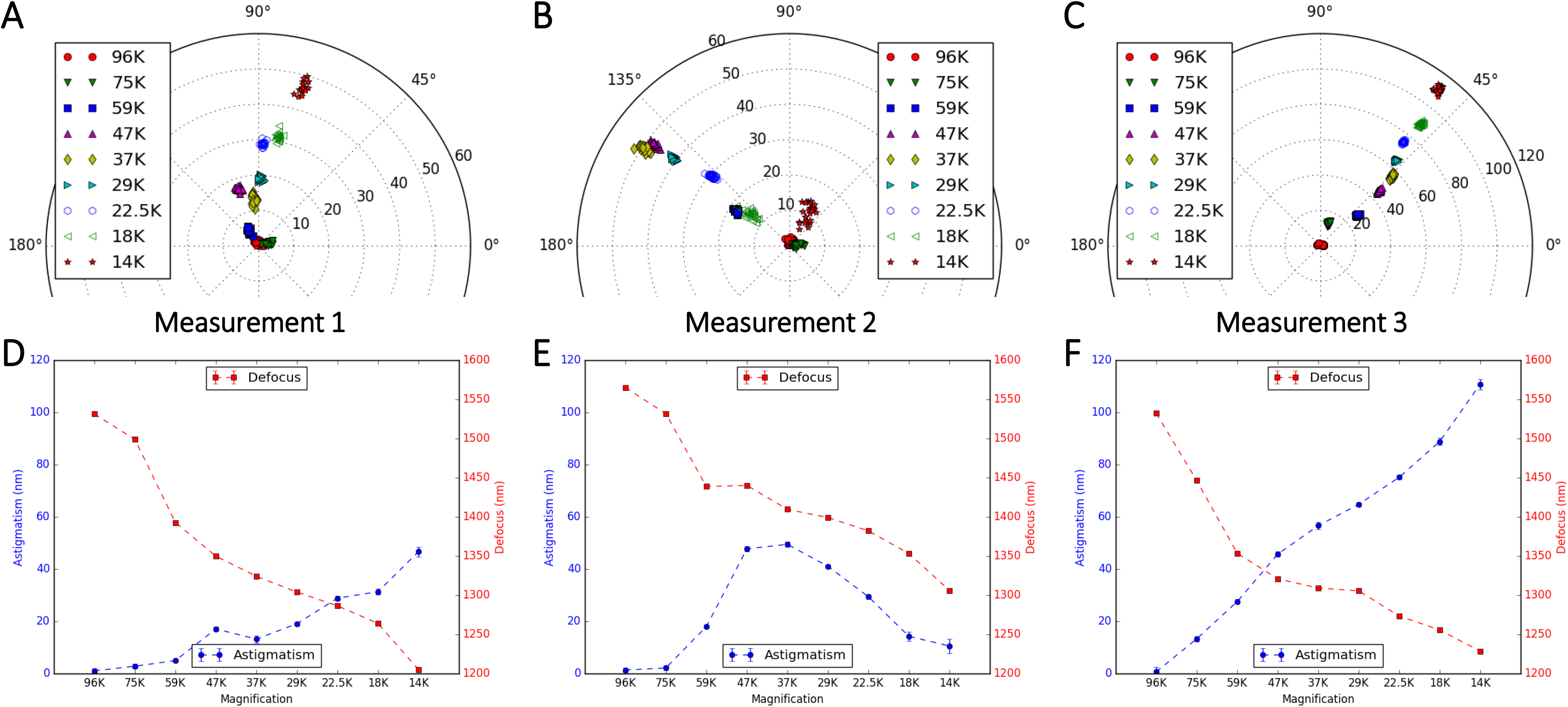
Magnification-dependent astigmatism detected on Titan Krios microscope. (A-C) Plots of astigmatism polar distribution with stepwise reduction of magnifications in three repeated measurements in a single day when the astigmatism is corrected at a nominal magnification of 96,000X. (D-F) The profiles of the variations of astigmatism (blue line) or defocus (red line) with the change of magnifications, corresponding to the measurements in (A-C). At each magnification, twenty images were collected and their astigmatisms and defocuses were calculated. The point and error bar represent the mean and RMSD of astigmatism in the blue line or defocus in the red line, respectively. The error bar may be too small to be visible in the line plot.

**Fig. 6.**
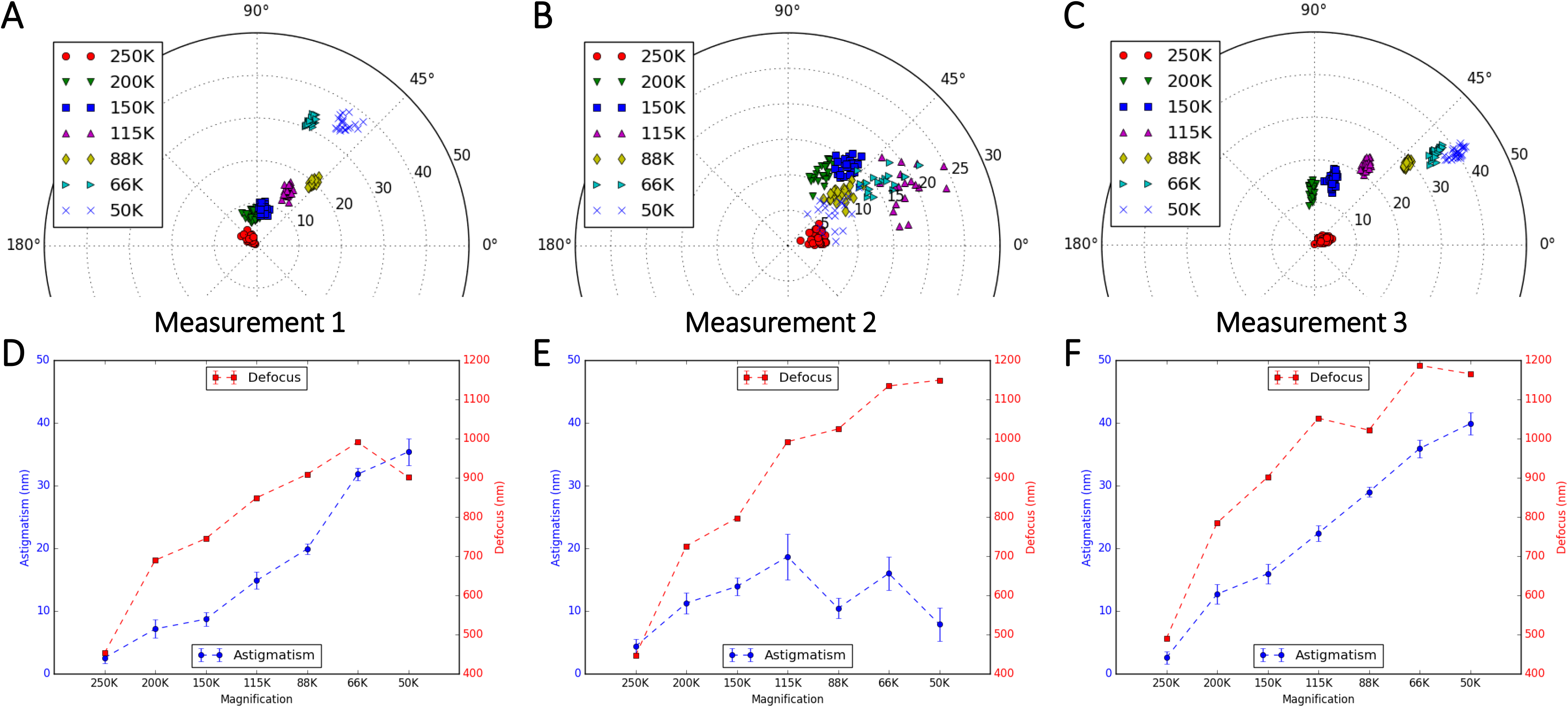
Magnification-dependent astigmatism detected on CM200 microscope. The measurements are the same as described in Fig. 5. The only difference being that the astigmatism is corrected at a nominal magnification of 250,000X on the CM200.

## 4. Discussion

In contrast to the dramatic progress in the automated cryo-EM data acquisition and image processing methods, little has changed for the microscope alignment tasks before data acquisition. In this paper, inspired by the observations of defocus-astigmatism correlations in experimental cryo-EM datasets, we have discovered the defocus/magnification-dependence of objective lens astigmatism and their stochastic variability using our recently published tool *s^2^stigmator*. These findings have essentially invalidated a basic assumption of current cryo-EM imaging strategy that assumes constant astigmatism for the significantly different defocuses/magnifications used in microscope alignment stage and final data acquisition stage.

### Vector summation model of the net astigmatism and the single pass tuning strategy

As shown here and in our previous work (Yan et al., 2017), *s^2^stigmator* can help achieve accurate correction of objective lens astigmatism at any imaging condition using a single pass tuning strategy. Understanding the principle of this single-pass tuning strategy will help further unveil the underlying theory of astigmatism variations with imaging parameters. Fig. 7 illustrates the vector diagrams (Fig. 7B-D) of three key points in a typical trajectory (Fig. 7A), including the initial point (? in Fig. 7A), the optimal point (➀ in Fig. 7A) in the arc-like segment when tuning the first stigmator (e.g. MY), and the final point at the center (➁ in Fig. 7A) after tuning the other stigmator (e.g. MX). The corresponding stigmator MX/MY values are labelled in parentheses next to the circled numbers. In Fig. 7B, *V*⃗_obj_, *V*⃗_*MX*_^0^, *V*⃗_*MY*_^0^ represents the initial state of the astigmatism of the objective lens, and the correction field of the stigmator MX and MY, respectively. *V*⃗_*sum*_^0^ is the summation of these three vectors and corresponds to the point ? in Fig. 7A. Here *V*⃗_*obj*_ can be considered as a constant vector in a short period of time, e.g. during astigmatism correction, while the correction field *V*⃗_*MX*_ and *V*⃗_*MY*_ will be varied to cancel *V*⃗_*obj*_ in order to minimize astigmatism. However, the orientations of *V*⃗_*MX*_^0^ and *V*⃗_*MY*_^0^ are fixed and the angle between them is also fixed at 45°, which are determined by the design of the octupole objective lens stigmator (Hawkes, 2013; Rai-Choudhury, 1997) assembly containing two interdigitated quadruple stigmators with 45° offset. When the stigmators are adjusted, the *V*⃗_*MX*_ and *V*⃗_*MY*_ vectors will change lengths without turning. The astigmatism correction task is to find the optimal lengths for both stigmator vectors so that the sum of the two correction vectors will be exactly inverse of the objective astigmatism vector *V*⃗_*obj*_ (i.e. same length but opposite direction). In Fig. 7C, only the stigmator MY is adjusted (red line in Fig. 7C) until it reaches the optimal length (*V*⃗_*MY*_^1^) at which its vector sum with *V*⃗_*obj*_ (*V*⃗_*sum*_^1^) is along the direction of *V*⃗_*MX*_. In this process, the resulted points (i.e. the net astigmatism, or the sum of the three vectors) of the trajectory shown in Fig. 7A exhibit an arc-like segment. Here the stigmator MX does not change (*V*⃗_*MX*_^1^ = *V*⃗_*MX*_^0^) and *V*⃗_*sum*_^1^ is along the direction *V*⃗_*MX*_^1^, corresponding to the point ➀ in Fig. 7A. In Fig. 7D, only the stigmator MX is adjusted to cause its correction field vector *V*⃗_*MX*_ to change length (green line in Fig. 7D) until the overall summation of vectors is 0 (*V*⃗_*sum*_^2^ = 0, ➁ in Fig. 7A). In this part of trajectory, the resulted points should directly move to the origin, forming a straight trace segment. As demonstrated in Fig. 7, the orientation of the straight trace segment is determined by the manufacturer's setting of the stigmator MX's direction. This finding can explain why the orientations of trajectories shown in Fig. S1 are the same, independent of defocus and magnification, when the two stigmators are adjusted in the order of MY first then MX. On the contrary, if the order of stigmators is switched during adjustment (MX first, then MY), the trajectory will rotate 45° and the straight trace segment will represent the direction of stigmator MY (Fig. 2A). This vector summation model is also validated on CM200 when the rotation of trajectories between different magnifications due to the imperfect rotation-free function is considered. The analysis described above explains the relationship of the objective lens astigmatism and the correction fields generated by the stigmators, and how the stigmators can be controlled to optimally compensate the objective lens astigmatism. The vector summation model described here further refines our previous model (Yan et al., 2017).

**Fig. 7.**
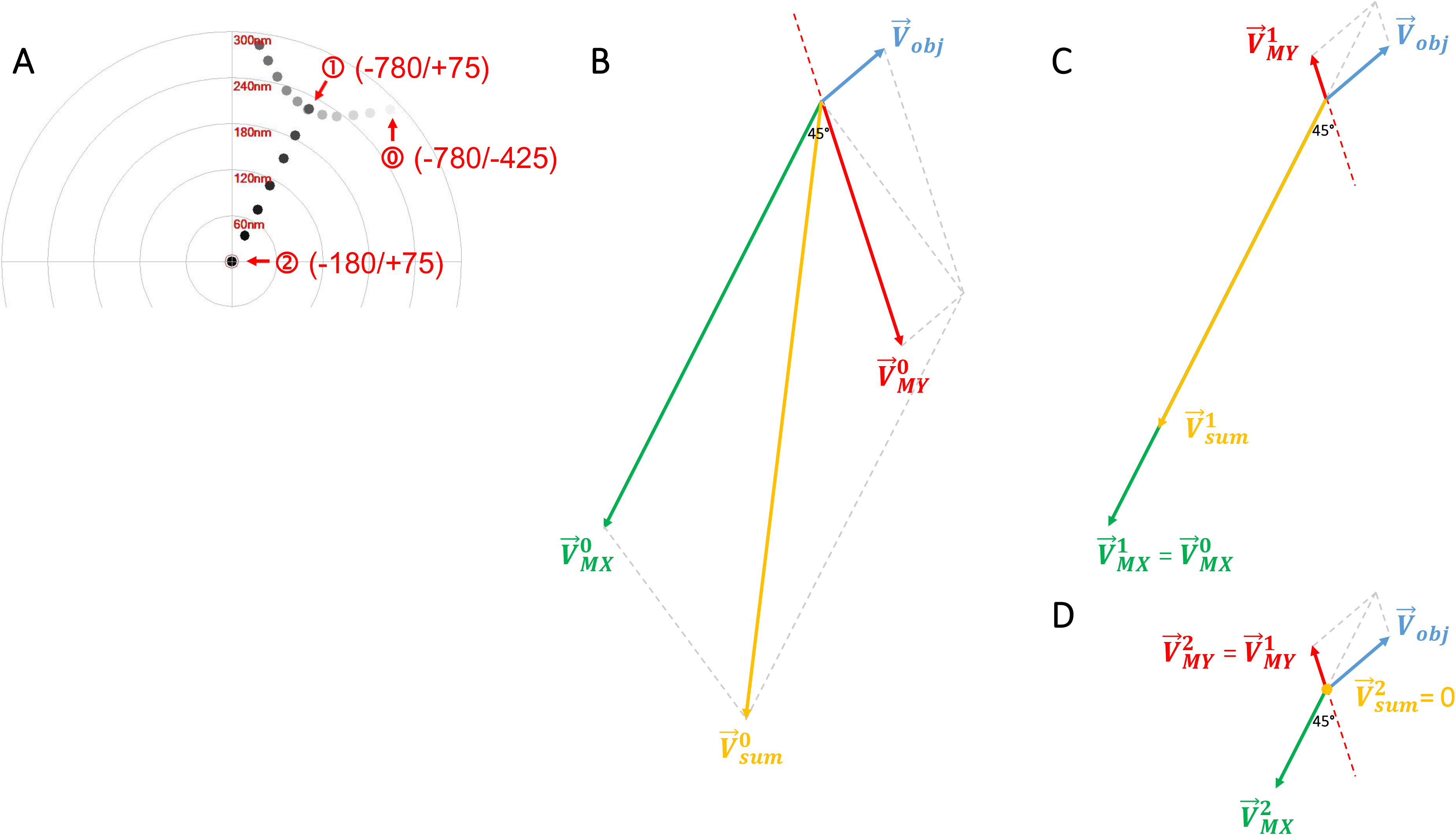
Vector diagrams to illustrate the principle of single-pass tuning strategy for astigmatism correction. (A) A screenshot of the trajectory on the Titan Krios microscope in which three key points are marked by red circled numbers ?, ?, ?, corresponding to the vector diagrams in (B-D), respectively. The corresponding stigmator MX/MY values are labelled in parentheses next to the circled numbers. (B) Initial point ?. The astigmatism of the objective lens, the correction fields of stigmator MX and MY are represented by *V*⃗_*obj*_, *V*⃗_*MX*_^0^, *V*⃗_*MY*_^0^, and their summation is represented by *V*⃗_*sum*_^0^. It is noted that *V*⃗_*obj*_ is assumed as a constant vector here, the directions of *V*⃗_*MX*_^0^ and *V*⃗_*MY*_^0^ are fixed and the angle between them is 45°. (C) The optimal point ➀ in the arc-like segment. The stigmator MY first reaches its optimal value (*V*⃗_*MY*_^1^) after adjusting along its own direction (red line) until *V*⃗_*sum*_^1^ is located in the direction of the stigmator MX (*V*⃗_*MX*_^1^). Here *V*⃗_*sum*_^1^ corresponds to the point ➀ in (A). (D) The final point ➁ of astigmatism correction. The stigmator MX is now adjusted along the direction of green line until *V*⃗_*sum*_^2^ is zero, corresponding to the point ➁ in (A). Consequently, the orientation of the straight trace segment in the trajectory (A) is determined by the orientation of stigmator MX.

### Defocus-dependent astigmatism

During single particle cryo-EM image acquisition, the defocuses for different images are intentionally varied to average out the effect of zero-nodes of CTF and obtain signals at all frequencies (Cheng et al., 2015; Penczek, 2010; Zhu et al., 1997). Nevertheless, the discovery of defocus-dependent astigmatism (Fig. 3) implies a significant problem of re-emerging astigmatism in this imaging strategy. Fig. 8 uses vector diagrams to explain the case of corrected astigmatism (Fig. 8A), and the re-emerged astigmatism after increasing (Fig. 8B) and decreasing (Fig. 8C) defocus. As can been seen from Fig. 8A, the total summation vector *V*⃗_*sum*_ = 0 when *V*⃗_*obj*_ is canceled by *V*⃗_*stigmator*_. Here *V*⃗_*obj*_ and *V*⃗_*stigmator*_ represent the astigmatism of objective lens and the combined correction field of the two stigmators, respectively. Moreover, the objective lens astigmatism (*V*⃗_*obj*_) is assumed to be proportional to the strength of objective lens current (*I*_*obj*_) as

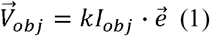

where *e⃗* is a unit vector representing the direction of the objective lens astigmatism (*V*⃗_*obj*_) and *k* is a scaling factor representing how strong the dependence is between *V*⃗_*obj*_ and *I*_*obj*_. In Fig. 8A, the astigmatism is corrected completely and the shape of Thon rings is perfectly circular (Fig. 8D). However, to increase defocus the objective lens current needs to be reduced to weaken the lens bending power, leading to a smaller objective lens astigmatism (blue arrow, *V*⃗_*df*↑_). The correction fields by the two stigmator (*V*⃗_*stigmator*_), which have not been changed from previous values optimized for a larger amount of objective lens astigmatism, now over-corrects the new, reduced objective lens astigmatism. A non-zero *V*⃗_*sum*_ (Fig. 8B) now appears and the Thon rings (Fig. 8E) become elongated. The analysis in Fig. 8B agrees with the observations from the Titan Krios (red line in Fig. 3A, Fig. 3B) and CM200 (red line in Fig. 3E, Fig. 3F) microscopes in the case of increasing defocus. Similarly, another non-zero *V*⃗_*sum*_ (Fig. 8C) appears in the opposite direction when defocus decreases, resulting in the Thon rings (Fig. 8F) becoming elongated along the perpendicular direction (blue arrow, *V*⃗_*df*↓_), since the switch of *V*⃗_*sum*_ direction is equivalent to change the ellipticity by 90°. The analysis in Fig. 8C also agrees with the observed defocus-dependence of astigmatism as defocus decreases (blue line in Fig. 3A, Fig. 3D for Titan Krios; blue line in Fig. 3E, Fig. 3H for CM200). Combining the vector diagrams in both Fig. 8B and C, the bi-directional increment of astigmatism can also be clearly understood, as well as the 90° angle between two branches in the polar plots (green line in Fig. 3A, Fig. 3C for Titan Krios; green line in Fig. 3E, Fig. 3G for CM200) when the astigmatism is minimized at the middle point of the defocus range. This finding of defocus-dependent astigmatism is also consistent with theoretic predictions based on Zernike polynomial expression of lens aberrations (Vargas et al., 2013).

**Fig. 8.**
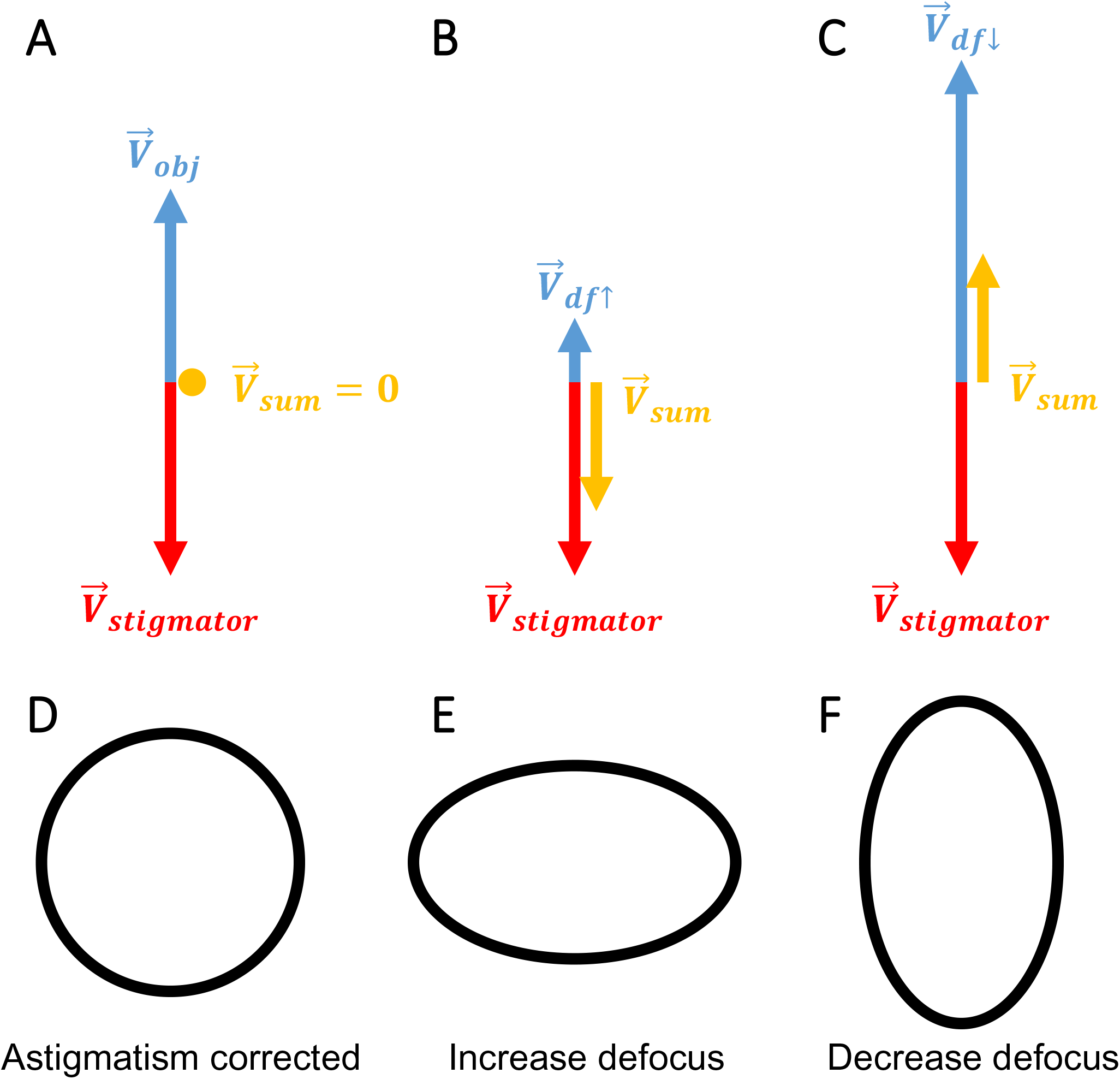
Vector diagram to interpret the defocus-dependent astigmatism shown in Fig. 3. (A-C) Vector diagrams illuminate the state of astigmatism fully corrected at a defocus (A), increased defocus after correction (B), decreased defocus after correction (C). Here the red arrows represent the combined correction field of the two stigmators (*V*⃗_*stigmator*_) and the blue arrows represent the astigmatism of the objective lens (*V*⃗_*obj*_) which is proportional to the strength of objective lens current. When defocus increases or decreases, the associated objective lens astigmatism, *V*⃗_*df*↑_ and *V*⃗_*df*↓_, becomes smaller or larger than the original *V*⃗_*obj*_ which is compensated by *V*⃗_*stigmator*_. (D-F) The representative sketches of the shape of Thon rings in the astigmatism states corresponding to the vector diagrams depicted in (A-C), respectively.

### Quantification of objective lens asymmetry

In the line plots from the Titan Krios (Fig. 3A) and CM200 (Fig. 3E), it is evident that the slopes of the linear trends are different for the two instruments. The slope measures how strong the dependence of astigmatism on defocus is and should be proportional to the scaling factor *k* in Eq. (1). For a perfectly round lens *k* is equal to 0 and as asymmetry in the lens increases, the larger *k* becomes. Therefore, we can use *k* as a parameter to quantify the quality of a TEM magnetic lens in terms of its cylindrical symmetry. A lens with smaller *k* will be a higher quality lens. Using this criterion, the objective lens of the Titan Krios microscope is more cylindric than that of CM200 microscope. This is consistent with the common understanding of current generation Titan Krios as a higher quality TEM than the CM200 microscope which was produced more than two decades ago. We propose that the defocus-dependent plots of astigmatism as shown in Fig. 3A and 3E are convenient measurements of the asymmetry level of the objective lens of a TEM instrument. Such quantitative measurements can be useful in several applications. For example, it can be used as one of the acceptance tests after the installation of a new TEM instrument. It can also be used to monitor the performance of the objective lens and to detect potential deterioration, for example, caused by a large contamination in the objective lens area.

### Stochastic variations of defocus-dependent astigmatism

The defocus-dependent astigmatism, including both the slope and the direction, was found to vary in our data (Fig. 4) when the same measurement was repeated after more than two-weeks. While the astigmatism of objective lens (*V*⃗_*obj*_) is considered stable within a short period time (a few hours to one or two days), it is also well-known that astigmatism tends to vary. As a result, it is a common practice to check and re-correct astigmatism as one of the daily-instrument alignment tasks. Our measurements (Fig. 4) have thus quantitatively verified the variability and validated the need for daily correction of astigmatism. Such variability can also be explained using the vector summation model (Fig. 9). *V*⃗_*obj*_ can vary due to the change of either the unit vector *e⃗* direction or the amplitude of the scaling factor (Eq. (1)) at different times. In the vector diagram (Fig. 9), the varying *V*⃗_*obj*_ (the objective lens astigmatism, blue arrows) is compensated by corresponding *V*⃗_*stigmator*_ (stigmator values, red arrows) that needs to be updated from day to day. There is a wide variety of reasons for the stochastic changes of *e⃗* and *k*, such as objective lens asymmetry due to imperfect manufacturing processes, electronic instability of the voltage and power supplies, column contaminations, temperature fluctuations of the objective lens chilling water, *etc*. (Barthel and Thust, 2013).

**Fig. 9.**
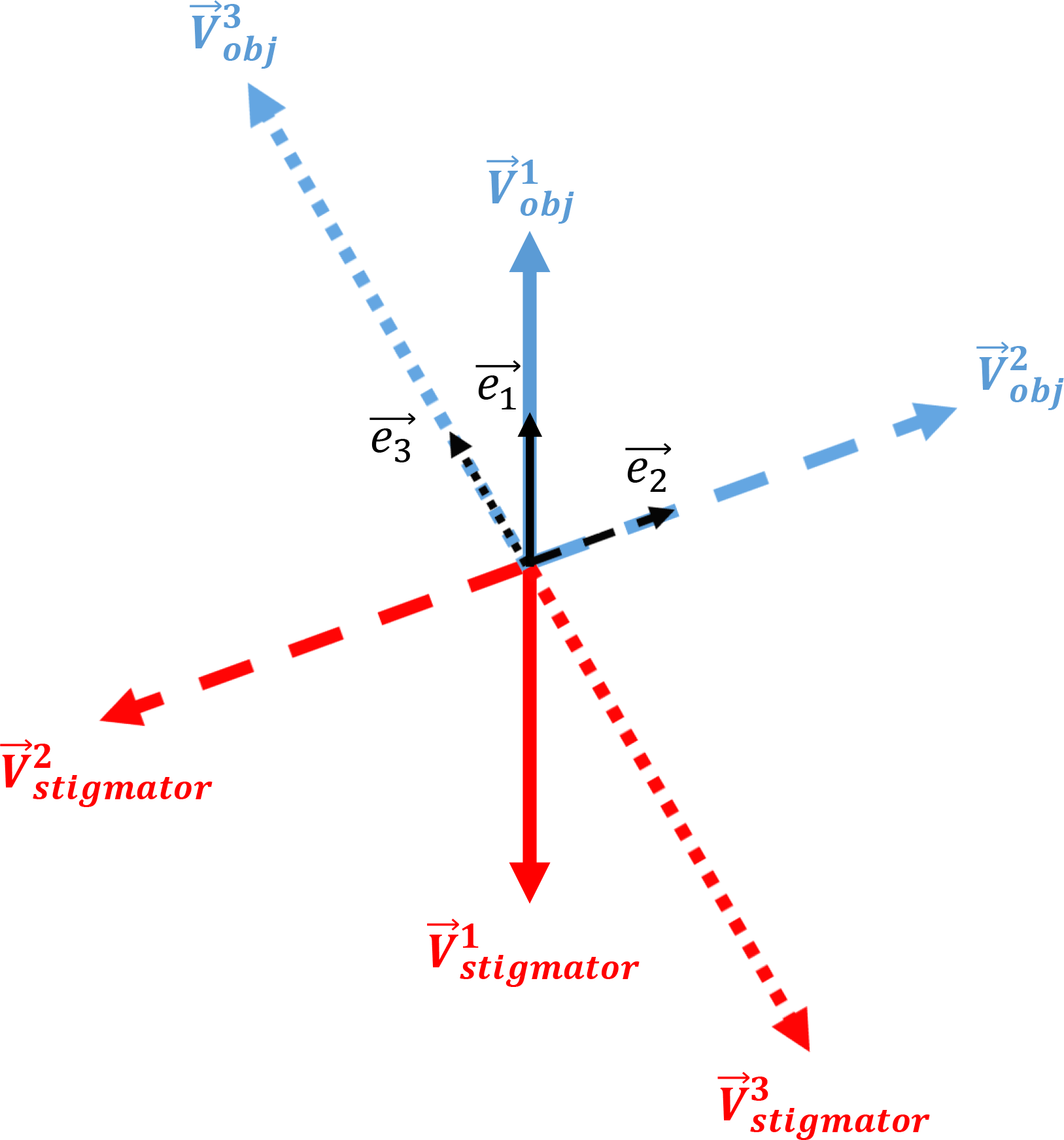
Vector diagram to interpret the variability of defocus-dependent astigmatism shown in Fig. 4. Here the vector representing the astigmatism of the objective lens (*V*⃗_*obj*_, blue arrows) varies from day to day. The randomness of *V*⃗_*obj*_ is determined by the orientation of the unit vector in black, *e⃗*_1_, *e⃗*_2_ and *e⃗*_3_, coupled with different scaling factors *k* (Eq. (1)). The combined effect of two stigmators (*V*⃗_*stigmator*_), represented by red arrows, needs to be varied accordingly to cancel *V*⃗_*obj*_ on different days.

### Magnification-dependent astigmatism

In addition to the defocus-dependent astigmatism, magnification-dependent astigmatism was also observed (Figs. 5 and 6). This implies that noticeable astigmatism would re-emerge during data collection if a different magnification is used for the correction of astigmatism during instrument alignment. Compared with the variability of astigmatism due to defocus, the astigmatism dependence on magnification is even more variable. When the same test was repeated three times on Titan Krios on the same day, the distribution of astigmatism at different magnifications is considerably different in both amplitude and angle (Fig. 5A-C). In the line plot for each measurement, the profile of astigmatism variation appears random in different tests (blue lines in Fig. 5D-F) while the profile of defocus variation is much more reproducible. Similar observations were obtained for both Titan Krios (Fig. 5) and CM200 (Fig. 6), which indicates that the defocus change is stable but the astigmatism change is unpredictable.

Modern TEM instruments usually use multiple imaging lenses, including an objective lens, a diffraction lens, an intermediate lens, and two projector lenses, to provide a wide range of magnifications. The astigmatism measured in the TEM image is a combined result of the astigmatism of all these lenses. The two stigmators actually correct the combined astigmatism of all these imaging lenses. When the magnification is changed, the current of all or a subset of these lenses would change, which leads to the change of individual lens astigmatism (Eq. (1)) and the combined astigmatism. As the stigmators have been tuned to correct the combined astigmatism at a particular magnification, the change of magnification will thus lead to re-emerging of astigmatism in the image at a different magnification. Since the currents of these lenses need to be changed in a non-linear pattern to achieve rotation-free imaging at multiple total magnifications, the combined astigmatism thus also varies in a non-linear pattern (Figs. 5 and 6). The irreproducibility of the profile of magnification-dependent astigmatism are caused by some random factors, such as column contaminations. Since the change of any one of the five lenses will change the combined astigmatism, it is thus not surprising the irreproducibility of the profile of magnification-dependent astigmatism is significantly worse than the irreproducibility of the profile of defocus-dependent astigmatism that is only affected by a single lens, the objective lens. In contrast, the profile of magnification-dependent defocus is more reproducible than that of magnification-dependent astigmatism as the pattern of current change is the same and the focus length of the lenses is more resistant to the random factors affecting the lens astigmatism.

### Recommendations for optimal TEM operations

The astigmatism of TEM images has been shown here to vary with changes in imaging conditions (e.g. defocus, magnification), indicating that correction of astigmatism at high magnification and near-focus conditions by the current approach will not be optimal after switching to different conditions for data acquisition. What's worse, the dependence of astigmatism on the imaging conditions varies from time to time, so that astigmatism cannot be reliably compensated by pre-calibration of the instrument. Based on our systematic measurements and analyses in this work, we suggest that 1) the magnification used for instrument alignment should be the same as the one used for data collection; 2) the defocus used for correction of astigmatism during instrument alignment should be set at the median defocus of the defocus range intended for subsequent data collection; 3) the magnification used for the focus-mode in the search-focus-exposure iterations of low-dose imaging should be the same magnification that is used for the exposure mode. To optimally correct the astigmatism for all images, a fast, accurate, and automated method needs to be developed to avoid the defocus-dependent astigmatism by adaptively correcting the astigmatism at all focuses.

## Acknowledgements

This work was supported in part by NIH grant (1R01AI111095). We thank the Purdue Cryo-EM Facility (http://cryoem.bio.purdue.edu) for the use of the Titan Krios and CM200 microscopes. We thank Dr. Dinesh Yernool, Dr. Anoop Narayanan, and Mr. Frank Vago for the RNA polymerase dataset, and Ms. Brenda Gonzalez for her assistance in preparation of the manuscript.

---

**Fig. S1.**
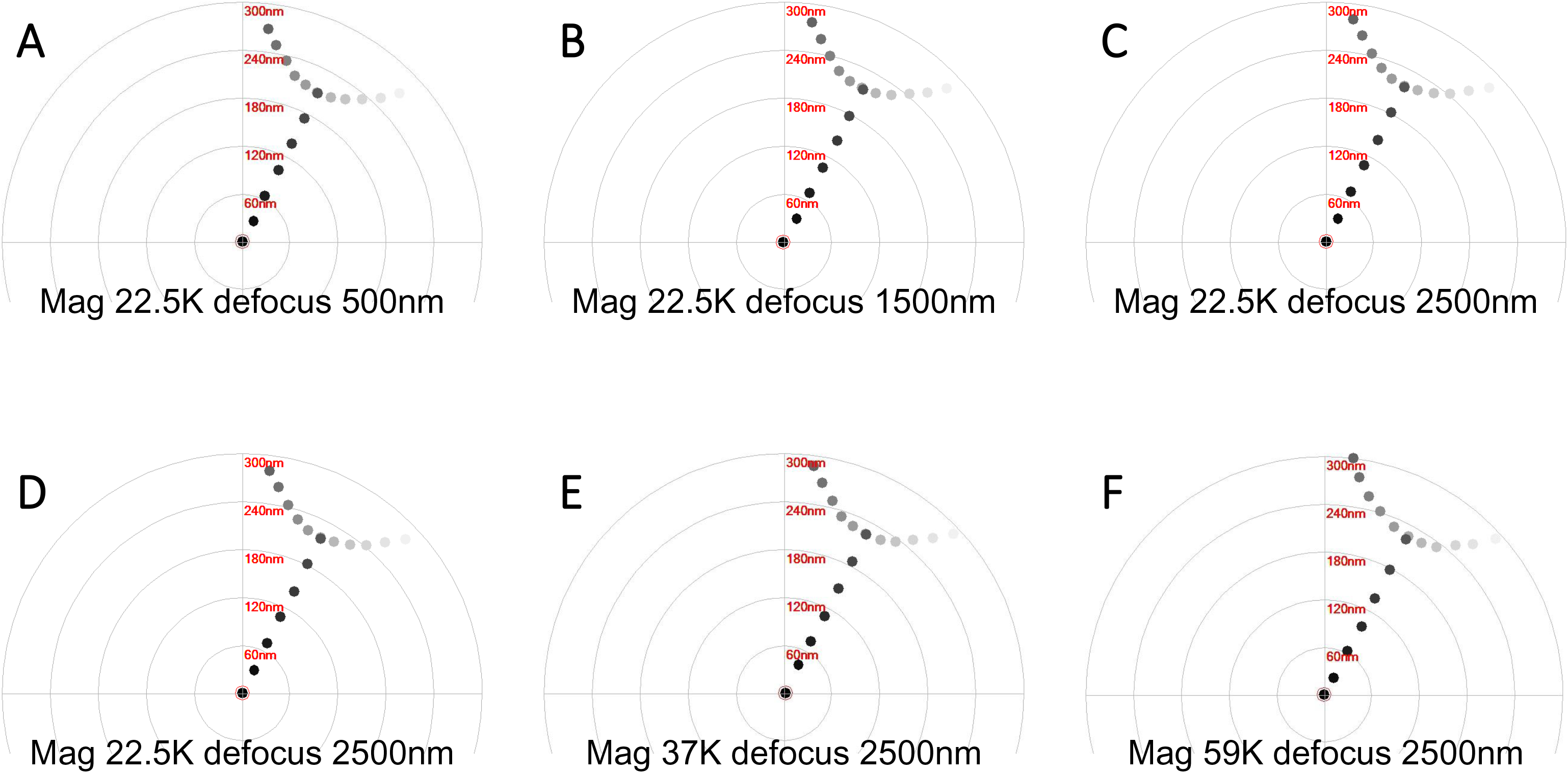
Representative trajectories of astigmatism correction at varying defocuses and magnifications on Titan Krios microscope. (A-C) The screenshots of the trajectories acquired at a nominal magnification of 22,500X and defocus 500 nm (A), 1500 nm (B) and 2500 nm (C), respectively. (D-F) The screenshots of the trajectories acquired at a nominal magnification of 22,500X (D), 37,000X (E) and 59,000X (F), respectively, and defocus 2500 nm.

**Fig. S2.**
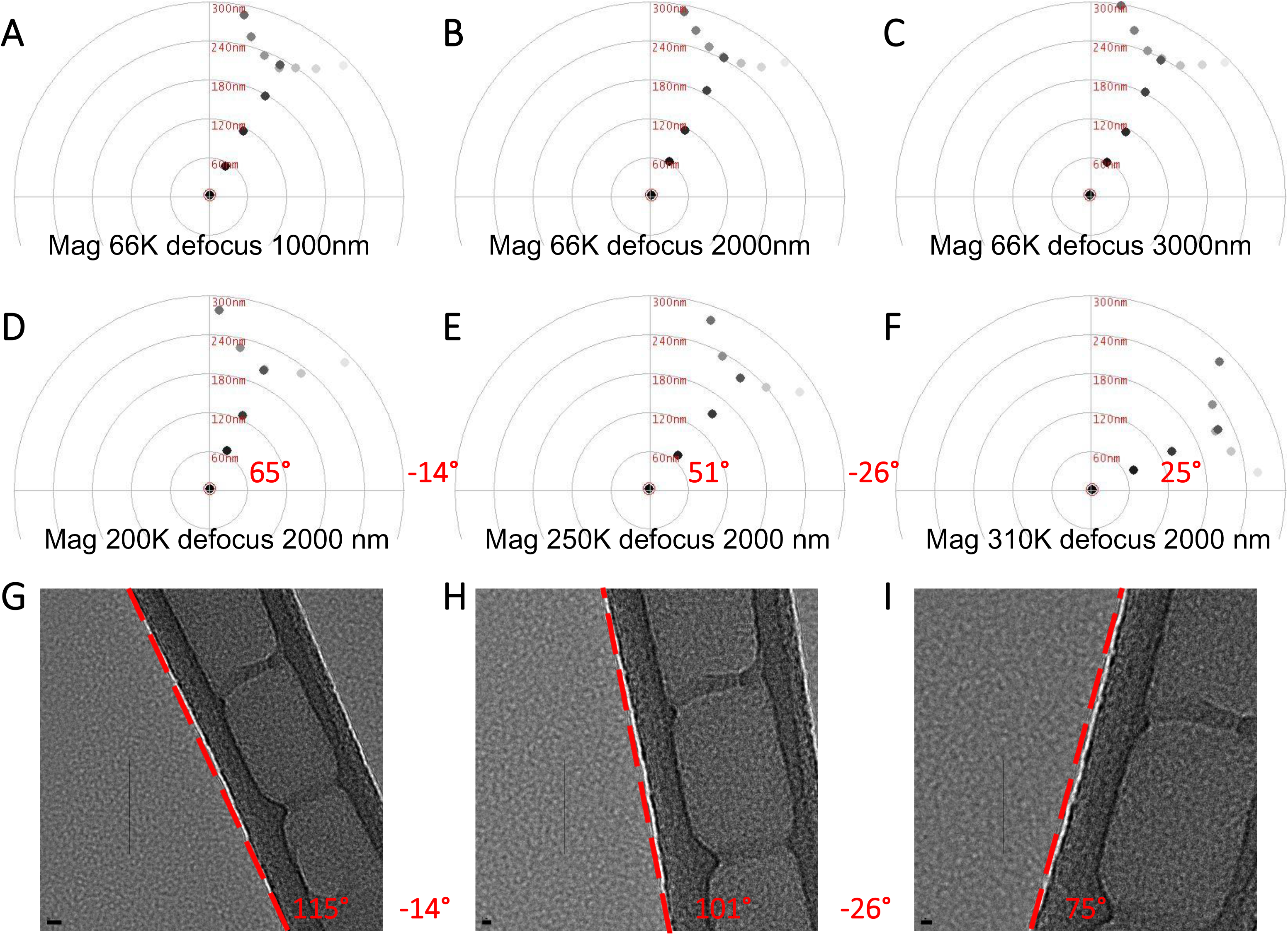
Representative trajectories of astigmatism correction at varying defocuses and magnifications on CM200 microscope. (A-C) The screenshots of the trajectories acquired at a nominal magnification of 66,000X and defocus 1000 nm (A), 2000 nm (B) and 3000 nm (C), respectively. (D-F) The screenshots of the trajectories acquired at a nominal magnification of 200,000X (D), 250,000X (E) and 310,000X (F), respectively, and defocus 2000 nm. The angles of the straight trace segments are 65°, 51° and 25°, respectively. And the difference between two adjacent trajectories are 14° and 26°. (G-I) The real space images collected at a nominal magnification of 200,000X (G), 250,000X (H) and 310,000X (I), respectively. The rotation of the red dash line represents the rotation of the image in real space with the change of magnifications. The angles of the red dash lines are 115°, 101° and 75°, respectively. And the angular difference between two adjacent images are 14° and 26°, in agreement with those of the trajectories shown in (D-F).

